# South Asia as a reservoir for the global spread of ciprofloxacin resistant *Shigella sonnei*

**DOI:** 10.1101/041327

**Authors:** Hao Chung The, Maia A. Rabaa, Duy Pham Thanh, Niall De Lappe, Martin Cormican, Mary Valcanis, Benjamin P. Howden, Sonam Wangchuk, Ladaporn Bodhidatta, Carl Jeffries Mason, To Nguyen Nguyen Thi, Duong Vu Thuy, Corinne N. Thompson, Nguyen Phu Huong Lan, Phat Voong Vinh, Tuyen Ha Thanh, Paul Turner, Poda Sar, Guy Thwaites, Nicholas R. Thompson, Kathryn E. Holt, Stephen Baker

## Abstract

**Background:** Antimicrobial resistance is a major issue in the *Shigellae*, particularly as a specific multidrug resistant (MDR) lineage of *Shigella sonnei* (lineage III) is becoming globally dominant. Ciprofloxacin is a recommended treatment for *Shigella* infections. However, ciprofloxacin resistant *S. sonnei* are being increasingly isolated in Asia, and sporadically reported on other continents.

**Methods and Findings:** Hypothesising that Asia is the hub for the recent international spread of ciprofloxacin resistant *S. sonnei*, we performed whole genome sequencing on a collection of contemporaneous ciprofloxacin resistant *S. sonnei* isolated in six countries from within and outside of Asia. We reconstructed the recent evolutionary history of these organisms and combined these data with their geographical location of isolation. Placing these sequences into a global phylogeny we found that all ciprofloxacin resistant *S. sonnei* formed a single clade within a Central Asian expansion of Lineage III. Further, our data show that resistance to ciprofloxacin within *S. sonnei* can be globally attributed to a single clonal emergence event, encompassing sequential *gyrA*-S83L, *parC*-S80I and *gyrA*-D87G mutations. Geographical data predict that South Asia is the likely primary source of these organisms, which are being regularly exported across Asia and intercontinentally into Australia, the USA and Europe.

**Conclusions:** This study shows that a single clone, which is widespread in South Asia, is driving the current intercontinental surge of ciprofloxacin resistant *S. sonnei* and is capable of establishing endemic transmission in new locations. Despite being limited in geographical scope, our work has major implications for understanding the international transfer of antimicrobial resistant *S. sonnei*, and provides a tractable model for studying how antimicrobial resistant Gram-negative community acquired pathogens spread globally.

## Introduction

Diarrheal disease is the second most common cause of mortality in children under the age of five years worldwide, equating to approximately 800,000 deaths per year [1]. The recent Global Enteric Multicentre Study (GEMS), a large prospective case-control study focusing on mild and severe paediatric diarrheal illnesses in sub-Saharan Africa and South Asia, found that *Shigella* (a genus of Gram-negative enteric bacteria) were amongst the top four most prevalent diarrhoeal pathogens in these settings [2]. The most recent estimates suggest that *Shigella* infections account for around 125 million cases of diarrhoea annually, with the majority occurring in children in low income countries [3]. There are four *Shigella* species (*dysenteriae*, *boydii*, *flexneri* and *sonnei*), but the overwhelming majority of the current global burden is presently caused by *S. sonnei* and *S. flexneri*. Present-day international epidemiology of the various *Shigella* species is particularly intriguing, as *S. sonnei* is replacing *S. flexneri* as the most common cause of shigellosis worldwide; this pattern is accentuated in regions undergoing rapid economic development [4,5], where *S. flexneri* dominated as recently as a decade ago.

*Shigella* infections are characterised by the invasion and disruption of the epithelial cells lining the gastrointestinal mucosa, resulting in mucous and/or bloody diarrhoeal discharge. Although shigellosis is typically self-limiting, antimicrobial treatment is used to prevent complications, reduce dysenteric discharge and curb post-symptomatic faecal shedding [6,7]. Consequently, resistance to antimicrobials restricts treatment options, placing vulnerable individuals suffering from shigellosis at increased risk of complications and increasing the likelihood of protracted faecal shedding. One of the current recommended first-line treatments for shigellosis is the fluoroquinolone, ciprofloxacin [8]. The fluoroquinolones target the DNA gyrase, a type II topoisomerase that is essential for bacterial DNA replication and transcription [9].

Antimicrobial resistance is an emerging global issue in *S. sonnei*, with a specific multidrug resistant (MDR) lineage (III) now dominating internationally. Further, organisms belonging to lineage III appear to be highly proficient at acquiring resistance to additional antimicrobials (including third generation cephalosporins) when they are introduced into new locations [10]. However, given their common usage and broad spectrum of activity, resistance against the fluoroquinolones is the most concerning. Since the first isolation of *S. sonnei* with reduced susceptibility to ciprofloxacin in Japan in 1993 [11], ciprofloxacin resistant *S. sonnei* have been increasingly reported throughout Asia [12-14]. Furthermore, public health laboratories in several non-Asian countries with low incidences of shigellosis have reported the isolation of ciprofloxacin resistant *S. sonnei*, often from individuals reporting recent travel to locations with a high risk of shigellosis [15-17].

Whole genome sequencing has proven to be the gold standard for tracking the international dissemination of clonal bacterial pathogens [18,19], and we have previously exploited this method to study the phylogenetic structure and spread of *S. sonnei* at both national and intercontinental levels [10,20]. Hypothesising that Asia was a hub for the recent international spread of ciprofloxacin resistant *S. sonnei*, we performed whole genome sequencing and phylogenetic characterisation of a collection of ciprofloxacin resistant *S. sonnei* isolated from within and outside Asia, aiming to explore the origins of this growing international epidemic.

## Methods

### Strain collection

Aiming to investigate the current international upsurge in ciprofloxacin resistant *S. sonnei* in detail, we gathered a collection of 60 contemporary ciprofloxacin resistant *S. sonnei* from six countries for whole genome sequencing. The isolates originated from Asian countries with a high incidence of shigellosis (Vietnam, n=11; Bhutan, n=12; Thailand, n=1; Cambodia, n=1), as well as isolates from countries with a low incidence of shigellosis (Australia, n=19; Ireland, n=16). Twelve additional ciprofloxacin susceptible *S. sonnei* sequences from these settings were also included for phylogenetic context. All strains were isolated independently between 2010 and 2015; details of the isolates used in this study are shown in Table 1.

**Table 1.** The origins of the *Shigella* isolates and sequences used in this study

| Country | Isolates in Central Asia clade (N) | Ciprofloxacin resistant isolates (N) | Study or institute origin | Patient group | Region of recent travel history (N) | Sequencing platform/Public database | Ciprofloxacin susceptibility |
| --- | --- | --- | --- | --- | --- | --- | --- |
| Bhutan | 12 | 12 | Diarrhoeal disease surveillance in JDWNRH<sup>a</sup>, Thimphu, Bhutan (AFRIMS) | Hospitalised children <5 years old | NA | Illumina HiSeq 2000 | Disk diffusion/E-te |
| Vietnam | 11 | 11 | Diarrhoeal disease surveillance in Ho Chi Minh City, Vietnam (OUCRU) | Hospitalised children < 5 years old | NA | Illumina MiSeq | Disk diffusion/E-te |
| Thailand | 8 | 1 | Diarrhoeal disease surveillance in Thailand (AFRIMS) | Hospitalised children <5 years old | NA | Illumina HiSeq 2000 | Disk diffusion |
| Cambodia | 1 | 1 | Diarrhoeal disease surveillance in Siem Reap, Cambodia (COMRU) | Hospitalised children <5 years old | NA | Illumina HiSeq 2000 | Disk diffusion |
| Ireland | 20 | 16 | National *Salmonella*, *Shigella*, and *L. monocytogenes* Reference Laboratory, Galway, Ireland | Primarily patients with recent travel history | India (9), Germany (1), Morocco (1), No travel (5), Unknown (4) | Illumina HiSeq 2000 | Broth microdilutio |
| Australia | 20 | 19 | Microbiological Diagnostic Unit Public Health Laboratory in Melbourne, Australia | Patients with recent travel history | India (15), Cambodia (3), Thailand (1), Southeast Asia (1). | Illumina NextSeq | Agar dilution |
| USA | 14 | 10 | Genome Trackr; Centre for Disease Control and Prevention, USA | Unknown | Unknown | Illumina MiSeq (BioProject PRJNA218110) | *in silico* assessment QRDR mutations |

<sup>a</sup> Jigme Dorji Wangchuk National Referral Hospital NA, Not applicable

The *S. sonnei* isolates from countries with a high incidence of shigellosis in Asia were collected through diarrhoeal disease surveillance in Bhutan and Thailand and as part of on-going, local IRB approved, hospital based studies in Ho Chi Minh City, Vietnam and Siem Reap, Cambodia [14]. The target patient group for these studies were/are generally hospitalised children aged less than five years residing in close proximity to the study centres. The ciprofloxacin resistant *S. sonnei* from countries with a low incidence of shigellosis (outside Asia) were collected and characterized by the National *Salmonella*, *Shigella* and *Listeria monocytogenes* Reference Laboratory, Galway, Ireland, and the Microbiological Diagnostic Unit Public Health Laboratory, Melbourne, Australia. These isolates were generally, but not exclusively, obtained from patients reporting recent travel to countries with a high incidence of shigellosis in Asia (Table 1). Susceptibility to ciprofloxacin was determined by either disk diffusion, E-test, agar dilution or broth microdilution, depending on the collaborating institution, and susceptibility breakpoints were interpreted according to the European Committee on Antimicrobial Susceptibility Testing (http://www.eucast.org/clinical_breakpoints). Namely, resistance was determined as strains with a zone of inhibition ≤15 mm (5 µg disc) and/or a Minimum Inhibitory Concentration (MIC) >2 µg/ml against ciprofloxacin; the various location specific methods and resulting data are described in Table 1.

### Genome sequencing and analysis

All isolated *S. sonnei* were sub-cultured and subjected to DNA extraction prior to whole genome sequencing on various Illumina platforms to produce pair-ended short read sequence; the specific sequencing system and the resulting public database numbers are shown in Table 1. We additionally included 14 *S. sonnei* sequences from organisms isolated in the USA and deposited in the GenomeTrackr Project (NCBI BioProject number PRJNA218110). All sequences were mapped to the *S. sonnei* Ss046 reference sequence (Accession number: NC_007384) using SMALT (version 0.7.4) and SNPs were called against the reference and filtered using SAMtools [21]. To contextualize all ciprofloxacin resistant *S. sonnei* within the global phylogeny, we appended our collection to include 133 publicly available sequences from a previous global analysis (accession ERP000182) [20]. Previously characterized mobile genetic elements and putative recombination (predicted using Gubbins) were removed [20], resulting in a gap-free alignment of 211 non-duplicate pseudo-whole genome sequences of 4,738 SNPs. A whole genome phylogeny was inferred from this alignment using RAxML v8.1.3 under the GTRGAMMA substitution model, and sufficient bootstrap replicates were determined automatically using the extended majority rule (MRE) bootstrap convergence criterion. In order to obtain a refined phylogenetic structure of the Central Asia clade, we applied the aforementioned approach to a set of 97 *S. sonnei* sequences (86 novel sequences and 11 historical sequences) belonging to this clade. This resulted in an alignment of 1,121 SNPs, which was used for phylogenetic inference.

## Results

### Fluoroquinolone resistant Shigella sonnei in a global context

We constructed a whole genome phylogeny of *S. sonnei*, incorporating sequences from 133 globally representative isolates and 86 novel isolates from Vietnam, Cambodia, Thailand, Bhutan, Australia, Ireland and the USA. The novel sequences included 60 from ciprofloxacin resistant (MIC >2 µg/ml) organism and 26 from ciprofloxacin susceptible organisms (or of unknown ciprofloxacin susceptibility isolated in the USA). The overall tree topology reflected the previously described global phylogenetic structure [20], confirming the presence of four distinct lineages (I, II, III and IV); lineage III was the most commonly represented and the most widely geographically distributed (Figure 1A). All ciprofloxacin resistant *S. sonnei* formed a single well-supported monophyletic clade within the Central Asian expansion of Lineage III (Central Asia III); an MDR group that is closely related but distinct from the Global III clade (Figure 1A and 1B).

**Figure 1.**
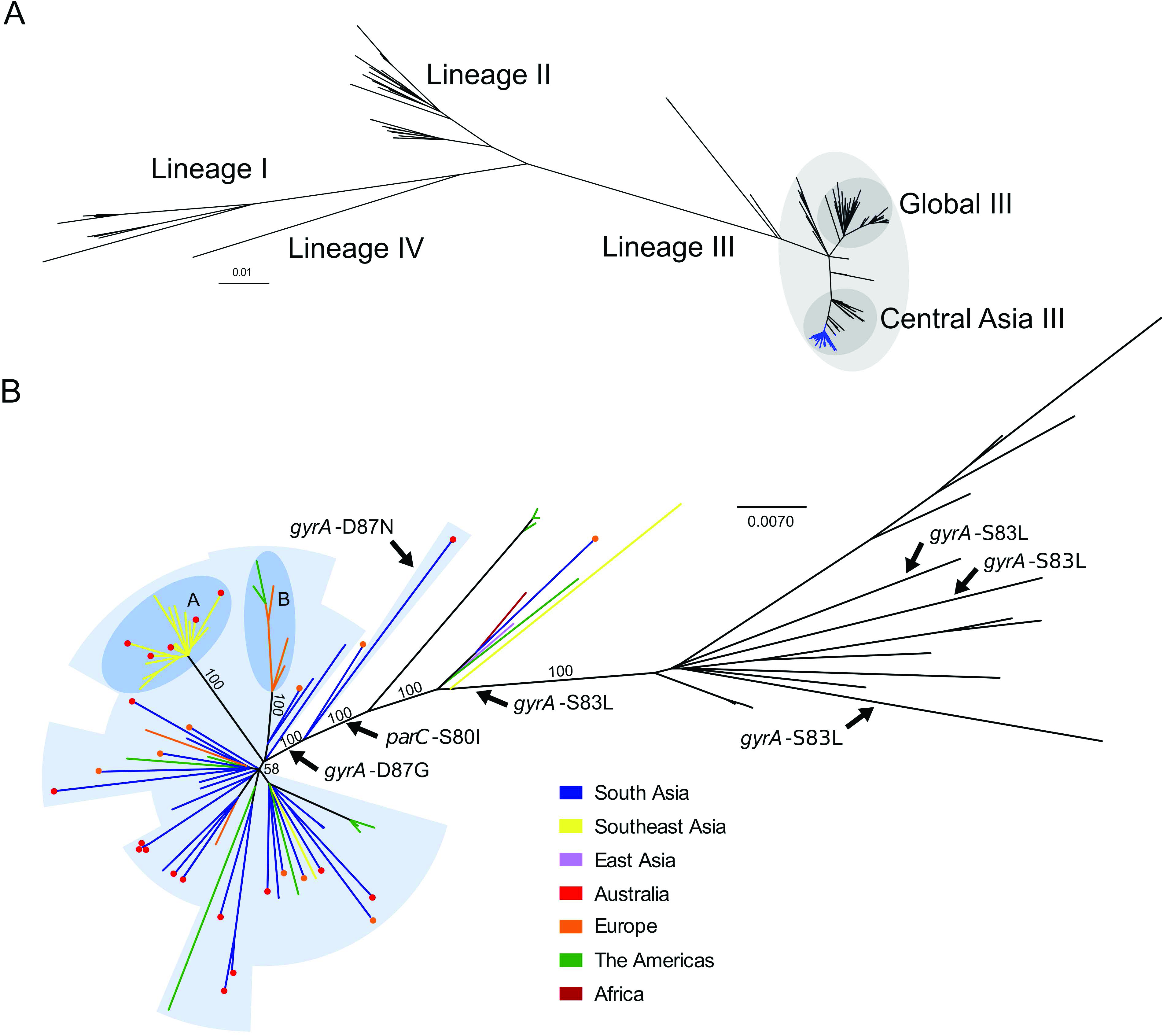
The phylogenetic structure of ciprofloxacin resistant *Shigella sonnei* in an international context. A) Unrooted maximum likelihood phylogeny of 211 globally representative *S. sonnei*, inclu sequences from ciprofloxacin resistant isolates (highlighted by the blue branches). Major li indicated by numbers (I, II, III and IV) as defined in Holt et al. 2012, with clades Global III Central Asia III within lineage III highlighted. Horizontal bar indicates the number of subst per site per year. B) Unrooted maximum likelihood phylogeny of Central Asia III, compris *sonnei* sequences. Branch colours indicate region of isolation (where no travel history is co region of recent travel (where travel history confirmed) according to the keys. For isolates with confirmed recent travel, a coloured circle at the tip indicates the region where the isolate w (multiple coloured circles are indicative of multiple isolates). Labelled arrows indicate bran where the mutations *gyrA*-S83L, *gyrA*-D87N, *gyrA*-D87G and *parC*-S80I have arisen. Blue background shading denotes isolates exhibiting ciprofloxacin resistance conferred by triple (*gyrA*-S83L, *parC*-S80I and *gyrA*-D87G (or *gyrA*-D87N)). Subpopulations A and B, are hi in the darker blue shaded areas, denoting clonal expansions in Southeast Asia and Europe/A respectively. Numbers above major branches indicate bootstrap support values, and horizon denotes the number of substitutions per site per year

### The emergence of a fluoroquinolone resistant Shigella sonnei clone

We next performed a more detailed phylogenetic reconstruction of the Central Asia III clade, incorporating sequence data from the 60 phenotypically ciprofloxacin resistant isolates and 26 others (ciprofloxacin susceptible or of unknown ciprofloxacin susceptibility), along with 11 historical Central Asia III sequences sourced from our previous global study (Figure 1B) [20]. The majority of the Central Asia III isolates carried more than three antimicrobial resistance genes, encoding resistance to a wide range of first-line drugs including tetracycline, streptomycin and co-trimoxazole. We additionally examined the genome sequence data for mutations in the Quinolone Resistance Determining Region (QRDR) within the DNA gyrase gene (*gyrA*) and the topoisomerase IV gene (*parC*), the regions encoding the target residues for fluoroquinolone activity. Overlaying these mutations on the phylogenetic tree indicated that the *gyrA*-S83L mutation, the first sequential mutation which confers reduced susceptibility against fluoroquinolones, has arisen independently within the Central Asia III clade on at least four separate occasions (Figure 1B). Amongst the isolates examined here for the first time, extensive resistance to ciprofloxacin can be attributed to a single clonal emergence event, via the sequential accumulation of *gyrA*-S83L followed by *parC*-S80I and *gyrA*-D87G, except for a single outlier that was isolated in Australia (Figure 1B). These three QRDR mutations were also shared by ten phenotypically uncharacterized *S. sonnei* from the USA, thus providing genotypic evidence for ciprofloxacin resistance. The single outlier isolate shares the *gyrA*-S83L and *parC*-S80I QRDR mutations of the other ciprofloxacin resistant isolates, but harbours *gyrA*-D87N rather than a *gyrA*-D87G, and is within a closely related out group of the major ciprofloxacin resistant clone (Figure 1B).

### South Asia as a hub of fluoroquinolone resistant Shigella sonnei

We additionally mapped the country of isolation and patient travel history onto the Central Asia III phylogeny to investigate the geographical structure of the clade (Figure 1B). For the ciprofloxacin resistant *S. sonnei* isolated from countries with a low incidence of shigellosis (Ireland, Australia and USA) and for which data on recent travel history was confirmed (27/45; 60%), India was the most commonly reported travel destination (21/27; 78%). The majority of the isolates associated with travel to India clustered closely with strains isolated in neighbouring Bhutan. These data suggest that South Asia is the primary source of ciprofloxacin resistant *S. sonnei* that have increasingly been isolated both inside and outside of Asia in recent years. Further, greater genetic diversity was observed within the South Asian *S. sonnei* than within the other sampled countries (Figure 1B), suggesting that this region acts as the most likely geographical source population.

Our data also show evidence of regional diversification of ciprofloxacin resistant *S. sonnei* within Asia. The phylogenetic structure is highly suggestive of a clonal expansion of ciprofloxacin resistant *S. sonnei* in Southeast Asia, specifically within Vietnam, as indicated by a long branch with 100% bootstrap support (Figure 1B). We additionally noted that *S. sonnei* nested within this clonal expansion were also isolated from travellers returning from countries including Cambodia and Thailand, indicating that isolates from this lineage have spread widely across Southeast Asia, as well as having been introduced into Australia on at least five separate occasions. An additional subpopulation of ciprofloxacin resistant *S. sonnei*, isolated in Ireland (five individuals with no recent history of travel and one individual returning from Germany) and the USA, are also likely representative of an expansion of this clone within Europe and the USA (Figure 1B). Whilst it was not possible to identify the geographical source definitively, the isolates most closely related to this European/USA subpopulation originated in India and Bhutan, again suggesting South Asia was the most likely origin. These two examples of subpopulation clonal expansions in Southeast Asia and Europe/USA indicate that this clone of ciprofloxacin resistant *S. sonnei* is also capable of sustained circulation upon introduction into new locations.

## Discussion

Here we provide direct evidence for the on-going global expansion of *S. sonnei* exhibiting new and clinically relevant antimicrobial resistance profiles. What is more, this study has significant implications for understanding the international trafficking of antimicrobial resistant bacterial pathogens. We suggest that as a single-serotype, human-adapted pathogen with a clonal population structure, *S. sonnei* serves as a tractable model for understanding how Gram-negative antimicrobial resistant pathogens are being regularly mobilised around the globe.

This is the first study that has used whole genome sequencing to examine the emergence and global spread of ciprofloxacin resistant *S. sonnei*. Our data show that all sequenced extant ciprofloxacin resistant *S. sonnei*, though sourced from disparate geographical locations, belonged to a single clonal expansion of Lineage III, with South Asia being the most likely hub for its origin and spread. Our findings support previous hypotheses suggesting that ciprofloxacin resistant *S. sonnei* in industrialised countries is being imported from South Asia [15,16]. A recent estimation of worldwide antimicrobial usage reported that India was the largest consumer of antimicrobials in 2010 [22]. Additionally, the fluoroquinolones are ranked as the most common antimicrobial prescribed for acute enteric diseases in India and Bangladesh [23,24]. The intensive use of fluoroquinolones in a region where there are foci of high population density and inconsistent access to sanitation is likely to have contributed to emergence of ciprofloxacin resistant enteric bacteria, such as *S. sonnei* and *Salmonella* Typhi, on the Indian subcontinent [19]. Global dissemination of these organisms is likely facilitated by the volume of travel between these regions and other areas of the world.

Our new data highlight the limitations of current typing protocols for tracking *S. sonnei*. It had been previously observed that some of the ciprofloxacin resistant *S. sonnei* isolates in this study (originating from Bhutan and Ireland) shared a similar *Xba*I Pulse Field Gel Electrophoresis (PFGE) pattern [14,15]. This pulsotype has been observed previously in India and Bangladesh [12,13,25-27], as well as in Canada [28], Belgium [29] and Japan [30], where the association with ciprofloxacin resistance was inconsistent. However, PFGE in this context did not offer sufficient granularity to link all of the isolates or provide sufficient resolution into the regional evolution of *S. sonnei*. Our phylogenetic analyses show that this pulsotype is associated with a phylogenetic lineage, supporting the notion that this pulsotype actually represents a widespread and pervasive subclade of Central Asia III.

This work has limitations. First, the lack of historical organisms from South Asia restricts our inference to only the contemporary situation. Further, additional present organisms from other settings would have improved our understanding of the current geographical spread of this clonal group. Notwithstanding these limitations, whole genome sequencing of these geographically disparate organisms has provided data at the highest resolution for deciphering the emergence and international spread of ciprofloxacin resistant *S. sonnei*. Future studies interrogating extensive spatial and temporal collections of ciprofloxacin resistant *S. sonnei*, as well as the *S. sonnei* diversity specific to South Asia prior to and during the emergence of antimicrobial resistance, are essential to further elucidate the origins and epidemiological dynamics of these populations. These supplementary investigations will greatly aid our efforts in controlling the spread of the current ciprofloxacin resistant clone and to prevent future emergent antimicrobial resistant bacterial populations.

In conclusion, the international surge of ciprofloxacin resistant *S. sonnei* clone poses a substantial global health challenge, and our data show this threat is not only manifested in sporadic cases from returning travellers but also the establishment of endemic transmission in new settings. The latter is already evident in high shigellosis incidence areas such as Southeast Asia. Therefore, integrative efforts from both the research community and public health authorities should be prioritised to track, monitor and prevent the international spread of this key enteric pathogen.

## Acknowledgements

We wish to acknowledge the staff of the Department of Enteric Disease, AFRIMS, Bangkok, Thailand for their work in isolating and maintaining the strains in this study. SB is a Sir Henry Dale Fellow, jointly funded by the Wellcome Trust and the Royal Society (100087/Z/12/Z). NRT is supported the Wellcome Trust grant #098051 to Wellcome Trust Sanger Institute. KEH is supported by fellowship #1061409 from the NHMRC of Australia. BPH is supported by fellowship #1023526 from the NHMRC of Australia. The funder of the study had no role in study design, data collection, data analysis, data interpretation, or writing of the report. The corresponding author had full access to all the data in the study and had final responsibility for the decision to submit for publication.

## References

1. Liu L, Johnson HL, Cousens S, Perin J, Scott S, Lawn JE, et al. Global, regional, and national causes of child mortality: an updated systematic analysis for 2010 with time trends since 2000. Lancet. Elsevier Ltd; 2012;379: 2151–61.

2. Kotloff KL, Nataro JP, Blackwelder WC, Nasrin D, Farag TH, Panchalingam S, et al. Burden and aetiology of diarrhoeal disease in infants and young children in developing countries (the Global Enteric Multicenter Study, GEMS): a prospective, case-control study. Lancet. 2013;382: 209–22.

3. Bardhan P, Faruque a SG, Naheed A, Sack D a. Decrease in shigellosis-related deaths without Shigella spp.-specific interventions, Asia. Emerg Infect Dis. 2010;16: 1718–23.

4. Vinh H, Nhu NT, Nga T V, Duy PT, Campbell JI, Hoang N V, et al. A changing picture of shigellosis in southern Vietnam: shifting species dominance, antimicrobial susceptibility and clinical presentation. BMC Infect Dis. 2009/12/17 ed. 2009;9: 204.

5. Thompson CN, Duy PT, Baker S. The Rising Dominance of Shigella sonnei: An Intercontinental Shift in the Etiology of Bacillary Dysentery. PLoS Negl Trop Dis. 2015;9: 1–13.

6. Bennish ML. Potentially lethal complications of shigellosis. Rev Infect Dis. 1991;13 Suppl 4: S319–S324.

7. Vinh H, Main J, Chinh M, Tam C, Trang P, Nga D, et al. Treatment of bacillary dysentery in Vietnamese children: two doses of ofloxacin versus 5-days nalidixic acid. Trans R Soc Trop Med Hyg. 2000;94: 323–326.

8. Guidelines for the control of shigellosis, including epidemics due to Shigella dysenteriae type 1. World Health Organization. 2005.

9. Ruiz J. Mechanisms of resistance to quinolones: target alterations, decreased accumulation and DNA gyrase protection. J Antimicrob Chemother. 2003; 1109–1117.

10. Holt KE, Vu T, Nga T, Pham D, Vinh H, Wook D, et al. Tracking the establishment of local endemic populations of an emergent enteric pathogen. PNAS. 2013;110.

11. Horiuchi S, Inagaki Y, Yamamoto N, Okamura N, Imagawa Y, Nakaya R. Reduced susceptibilities of Shigella sonnei strains isolated from patients with dysentery to fluoroquinolones. Antimicrob Agents Chemother. 1993;37: 2486–2489.

12. Nandy S, Dutta S, Ghosh S, Ganai A, Rajahamsan J, Theodore RBJ, et al. Foodborneassociated Shigella sonnei, India, 2009 and 2010. Emerg Infect Dis. 2011;17: 2072–2073.

13. Ud-Din AIMS, Wahid SUH, Latif H a, Shahnaij M, Akter M, Azmi IJ, et al. Changing trends in the prevalence of Shigella species: emergence of multi-drug resistant Shigella sonnei biotype g in Bangladesh. PLoS One. 2013;8: e82601.

14. Ruekit S, Wangchuk S, Dorji T, Tshering KP, Pootong P, Nobthai P, et al. Molecular characterization and PCR-based replicon typing of multidrug resistant Shigella sonnei isolates from an outbreak in Thimphu, Bhutan. BMC Res Notes. 2014;7: 95.

15. De Lappe N, Connor JO, Garvey P, Mckeown P, Cormican M. Ciprofloxacin-Resistant Shigella sonnei Associated with Travel to India. Emerg Infect Dis. 2015;21: 2013–2015.

16. Bowen A, Hurd J, Hoover C, Khachadourian Y, Traphagen E, Harvey E, et al. Importation and Domestic Transmission of Shigella sonnei Resistant to Ciprofloxacin - United States, May 2014-February 2015. Morb Mortal Wkly Rep. 2015;64: 318–320.

17. Kim JS, Kim JJ, Kim SJ, Jeon S, Seo KY, Choi J, et al. Shigella sonnei Associated with Travel to Vietnam, Republic of Korea. Emerg Infect Dis. 2015;21: 1247–1250.

18. He M, Miyajima F, Roberts P, Ellison L, Pickard DJ, Martin MJ, et al. Emergence and global spread of epidemic healthcare-associated Clostridium difficile. Nat Genet. 2013;45: 109–13. doi:10.1038/ng.2478

19. Wong VK, Baker S, Pickard DJ, Parkhill J, Page AJ, Feasey NA, et al. Phylogeographical analysis of the dominant multidrug-resistant H58 clade of Salmonella Typhi identifies inter-and intracontinental transmission events. Nat Genet. 2015;47: 632–9. doi:10.1038/ng.3281

20. Holt KE, Baker S, Weill FX, Holmes EC, Kitchen A, Yu J, et al. Shigella sonnei genome sequencing and phylogenetic analysis indicate recent global dissemination from Europe. Nat Genet. 2012/08/07 ed. 2012;44: 1056–1059.

21. Li H, Handsaker B, Wysoker A, Fennell T, Ruan J, Homer N, et al. The Sequence Alignment/Map format and SAMtools. Bioinformatics. 2009;25: 2078–9.

22. Van Boeckel TP, Gandra S, Ashok A, Caudron Q, Grenfell BT, Levin S a., et al. Global antibiotic consumption 2000 to 2010: An analysis of national pharmaceutical sales data. Lancet Infect Dis. Elsevier Ltd; 2014;14: 742–750.

23. Fahad BM, Matin A, Shill MC, Asish KD. Antibiotic usage at a primary health care unit in Bangladesh. Australas Med J. 2010;3: 414–421.

24. Kotwani A, Chaudhury RR, Holloway K. Antibiotic-prescribing practice of primary care prescribers for acute diarrhea in New Delhi, India. Value Heal. 2012;15: S16–S119.

25. Pazhani GP, Niyogi SK, Singh AK, Sen B, Taneja N, Kundu M, et al. Molecular characterization of multidrug-resistant Shigella species isolated from epidemic and endemic cases of shigellosis in India. J Med Microbiol. 2008; 856–863.

26. Ghosh S, Pazhani GP, Chowdhury G, Guin S, Dutta S, Rajendran K, et al. Genetic characteristics and changing antimicrobial resistance among Shigella spp. isolated from hospitalized diarrhoeal patients in Kolkata, India. J Med Microbiol. 2011;60: 1460–1466.

27. Talukder K a., Islam Z, Dutta DK, Aminul Islam M, Khajanchi BK, Azmi IJ, et al. Antibiotic resistance and genetic diversity of Shigella sonnei isolated from patients with diarrhoea between 1999 and 2003 in Bangladesh. J Med Microbiol. 2006;55: 1257–1263.

28. Gaudreau C, Ratnayake R, Pilon P a, Gagnon S, Roger M, Lévesque S. CiprofloxacinResistant Shigella sonnei among Men Who Have Sex with Men, Canada, 2010. Emerg Infect Dis. 2011;17: 1747–1750.

29. Vrints M, Mairiaux E, Van Meervenne E, Collard JM, Bertrand S. Surveillance of antibiotic susceptibility patterns among Shigella sonnei strains isolated in Belgium during the 18-year period 1990 to 2007. J Clin Microbiol. 2009;47: 1379–1385.

30. Uchimura M, Kishida K, Koiwai K. Increasing Incidence and the Mechanism of Resistance of Nalidixic Acid Resistant Shigella sonnei. Kansenshogaku Zasshi. 2001;75: 923–930.

